# Basal leakage in oscillation: coupled transcriptional and translational control using feed-forward loops

**DOI:** 10.1101/2020.02.24.962415

**Authors:** Ignasius Joanito, Ching-Cher Sanders Yan, Jhih-Wei Chu, Shu-Hsing Wu, Chao-Ping Hsu

## Abstract

The circadian clock is a complex system that plays many important roles in most organisms. Previously, many mathematical models have been used to sharpen our understanding of the *Arabidopsis* clock. However, these models are mostly dependent on transcriptional regulation, and the importance of post-translational regulation is still rarely discussed from theoretical aspects. In this study, we built a series of simplified oscillators with different regulations to study the importance of post-translational regulation (specifically, 26S proteasome degradation) in the clock system. We found that a simple transcriptional-based oscillator can already generate sustained oscillation, but the oscillation can be easily destroyed in the presence of transcriptional leakage. Coupling post-translational control with transcriptional-based oscillator in a feed-forward loop will greatly improve the robustness of the oscillator in the presence of basal leakage. Using these general models, we were able to replicate the increased variability observed in the E3 ligase mutant for both plant and mammalian clocks. With this insight, we also predict a plausible regulator of several E3 ligase genes in the plant’s clock. Thus, our results provide insights into and the plausible importance in coupling transcription and post-translation controls in the clock system.

**Author summary:** For circadian clocks, several current models had successfully captured the essential dynamic behavior of the clock system mainly with transcriptional regulation. Previous studies have shown that the 26s (1, 2) proteasome degradation controls are important in maintaining the stability of circadian rhythms. However, how the loss-of-function or over-expression mutant of this targeted degradations lead to unstable oscillation is still unclear. In this work, we investigate the importance of coupled transcriptional and post-translational feedback loop in the circadian oscillator. With general models our study indicate that the unstable behavior of degradation mutants could be caused by the increase in the basal level of the clock genes. We found that coupling a non-linear degradation control into this transcriptional based oscillator using feed-forward loop improves the robustness of the oscillator. Using this finding, we further predict some plausible regulators of Arabidopsis’s E3 ligase protein such as COP1 and SINAT5. Hence, our results provide insights on the importance of coupling transcription and post-translation controls in the clock system.

---

The circadian clock is an endogenous time-keeping mechanism in cells that anticipates daily changes in the environment (3-6). It controls the daily rhythm of many biological processes (7-9) and disruption of the clock has been associated with many disadvantageous traits (10-13). Like many eukaryotes, in the *Arabidopsis* clock, the core oscillator is governed by coupled transcription and translation feedback loops (TTFL) (14). The transcriptional circuit consists of several important genes such as *CIRCADIAN CLOCK-ASSOCIATED1* (*CCA1*), *LATE ELONGATED HYPOCOTYL (LHY), TIMING OF CAB EXPRESSION 1 (TOC1), PSEUDO-RESPONSE REGULATOR 9 (PRR9), PRR7*, and *PRR5* (14). Experimentally, *CCA1* and *LHY* genes were found to repress the expression of *TOC1, PRR9, PRR7*, and *PRR5* (15-17), whereas in turn, all of them repressed *CCA1* and *LHY* expression (18-21). Furthermore, *TOC1* and *PRR5* can repress *PRR9* and *PRR7* expressions (19, 22). Together, they formed a 3-node loop of negative regulation, the repressilator (23). The core repressilator motif is coupled with positive feedback loops, leading to several interesting properties (24-26). Although current models have successfully helped us to understand the behavior of the *Arabidopsis* clock, they mostly still depend on the transcriptional regulation process, and the importance of post-translational regulations is still rarely discussed.

In *Arabidopsis*, many post-translational regulations, such as protein–protein interaction (27, 28), subnuclear localization (29), phosphorylation (30), and 26S proteasome degradation (1, 31), have been reported in the past decades. Among these many regulations, the 26S proteasome degradation pathway, also known as ubiquitin-proteasome system (UPS), attracts our attention. UPS involves in almost all aspects of plant life cycle, such as root elongation, light response, flowering time, seed development and also biotic and abiotic stress (for comprehensive review, see (32, 33)). Experimentally, almost all important clock genes, including *CCA1* (34), *LHY* (35, 36), *TOC1* (31), *PRR9* (37), *PRR7* (38), *PRR5* (39), *GIGANTEA* (*GI*) (40), *EARLY FLOWERING 3* (*ELF3*) and *CONSTITUTIVE PHOTOMORPHOGENIC* 1 (*COP1*) (41), were found degraded through the 26S proteasome degradation pathway. These observations imply that the 26s proteasome degradation pathway is important in the *Arabidopsis* clock system.

Remarkably, such degradation control is also found in other clock systems such as mammals, flies, and fungi (4-6, 42). In the mammalian clock, two important clock components, cryptochrome (*CRY*) and period (*PER*), are also degraded through the 26S proteasome degradation pathway. *CRY* protein is targeted for proteasomal degradation by two different F-box proteins, *Fbxl3* and *Fbxl21* (43-47). However, mutations in these two E3 ligase proteins did not alter the behavior of circadian rhythms markedly as compared with phenotypes that were caused by mutations in other clock genes (2). A recent study showed that alteration of *β-Trcp1* and *β-Trcp2* proteins, other F-box proteins that will trigger the degradation of *PER* protein, severely altered the clock’s function. In the *β-Trcp* double-mutant mice, the oscillation of many clock genes are highly unstable, as indicated by the greatly increased variability in circadian rhythm (2).

The striking increase in variability of E3 ligase mutant plant has also been seen in the *Arabidopsis* clock. Previously, an F-box protein, *ZEITUPLE (ZTL)*, was found involved in target degradation of both *PRR5* and *TOC1* (31, 39). Somers *et al.* showed that mutation of *ZTL* protein would greatly increase the variability of both amplitude and period of circadian oscillations (1). To the best of our knowledge, why this mutation can lead to high variability in the plant’s clock is still unknown. For the mammalian clock, D’Alessandro *et al.* proposed that such unstable circadian rhythms in *β-Trcp* mutant mice may come from loss of nonlinear degradation of *PER* protein (2). However, why such loss of nonlinear degradation can lead to unstable behavior is still not clear.

Hence, in this study, we developed a series of simplified models to study the importance of targeted degradation in the clock system. Our results showed that basal leakage in the transcription of clock genes leads to unstable behavior of the clock system, which can be stabilized by the degradation control. Here, we showed that basal leakage could easily reduce the robustness of an oscillator, especially for a pure transcriptional controlled oscillator. However, combining transcriptional and post-translational controls can greatly improve the robustness of the oscillator by providing another layer of regulation such that the system can still push the protein level back, despite leakage in the mRNA level. Moreover, we also found that coupling E3 ligase using a coherent feed-forward loop can give better control to the basal leakage as compared with other network motifs. Using these general models, we have successfully replicated the observed experimental results of the *ZTL* and *β-Trcp* mutant conditions and also predict plausible regulators of several E3 ligases in *Arabidopsis*. Therefore, our results provide plausible importance in coupling transcription and post-translation control in the clock system.

## Results

### Transcription-based oscillators have lower noise but are susceptible to transcriptional leakage

In this study, we built a series of simple oscillator systems based on the repressilator, with different regulation controls (Figure 1A). A repressilator is a circular three-inhibitor feedback loop that was originally constructed as a synthetic circuit and capable of generating oscillation in *Escherichia coli* (23). However, in recent years, repressilators can also be found in many oscillating systems such as *Arabidopsis* (48) and mammalian (49) circadian clocks. The system has also been widely used to study many interesting properties of an oscillator such as tunability (50) and switchability (51, 52). Therefore, we used a repressilator to study the effect of different regulation controls in the oscillator, including transcription (M1), post-translation (M2), transcription with positive auto-regulation (M3), post-translation with positive auto-regulation (M4), and combined transcription and post-translational control (M5) (Figure 1A). Here, we randomly selected the parameter sets for each model such that their deterministic dynamics would oscillate and then performed Gillespie’s stochastic simulation (53, 54). The robustness of the oscillation was measured by estimating the normalized decay rate of an autocorrelation function (Figure 1B, Methods).

**Figure 1.**
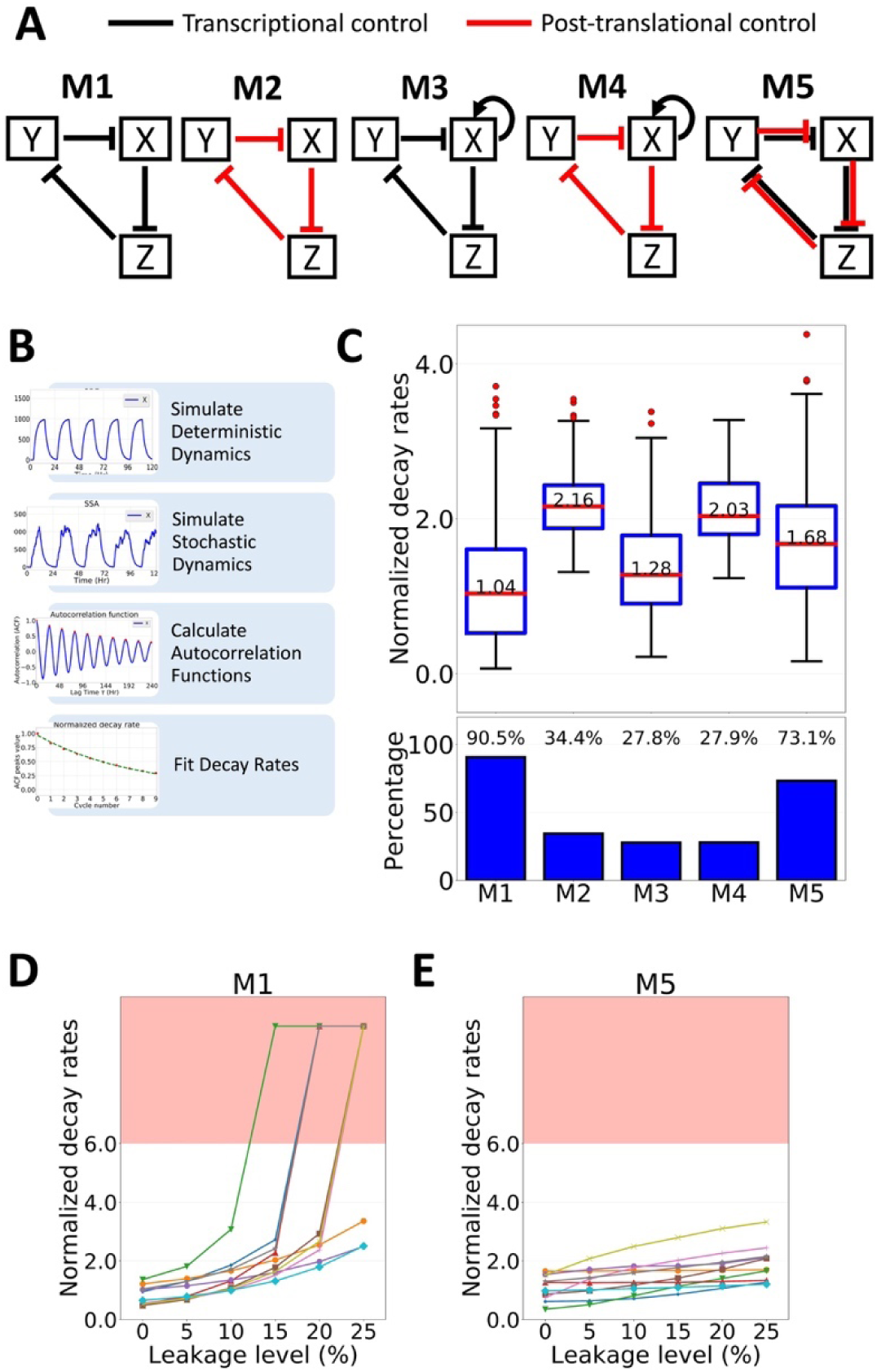
Combining transcriptional and post-translational controls improves the robustness of an oscillator. (A) Schematic representation of the tested models. (B) General workflow used in this study. (C, Upper panel) Box plot representing normalized decay rates of each tested model for 1000 randomly select parameter sets. Red lines indicate the median, and box edges indicate the 25th (Q1) and 75th (Q3) percentiles. Whiskers are plotted at 1.5*(Q3-Q1). (C, lower panel) Percentage of parameter sets that showed sustained oscillation under stochastic simulations. (D, E) the normalized decay rates of 10 randomly chosen parameter sets of M1 (D) or M5 (E) under different leakage levels. Each line represents one parameter set. The red-shaded region of the plot indicates non-oscillating results.

For a simple transcriptional feedback model, M1, among 1,000 parameter sets that can oscillate under deterministic propagation, 905 (90.5%) yielded sustained oscillation under a noisy condition, and their normalized decay rates have a median value of 1.04 (Figure 1C). For the post-translational feedback model, M2, oscillatory parameter sets under a noisy condition were found at a lower rate: 344 of 1,000 (34.4%), with much worse median value of 2.16 in decay rates (Figure 1C). This result may occur because, in M2, the noise from uncontrolled mRNA was propagated to the protein level, which easily altered the phase and period of the oscillation (Figure S1). Furthermore, adding an auto-positive feedback on a transcriptional-based oscillator (M3) or post-translational–based oscillator (M4) yielded oscillations at an even lower rate: 278 of 1,000 parameter sets (27.8%) with median value of 1.28 for M3, and 279 of 1,000 parameter sets (27.9%) with median value of 2.03 for M4. Thus, our results showed that a simple transcriptional-based oscillator is more robust than a post-translational-based oscillator.

Next, we tested the oscillators for transcriptional leakage, which commonly occurs in cells. Many studies have shown that promoters are actually leaky (55-58). Yanai *et al.* in 2006, even suggested that the selection against “unnecessary” transcription is low and hypothesized that leakiness of the promoters may be evolutionarily neutral (59, 60). Hence, we added an additional term to the gene production to represent this leakage in our models, and re-tested the performance of M1 (see methods for more detailed information). Surprisingly, adding transcriptional leakage greatly increased the noise level in model M1 (Figure 1D). Actually, the system was very sensitive such that adding 5% leakage reduced the number of oscillating parameters sets from 90.5% to 7.3% (Figure S2A). Furthermore, adding a positive feedback did not improve the performance of the transcriptional-based oscillator (Figure S2B). Therefore, these results suggested that although a simple transcriptional-based oscillator can already generate good oscillation, it is prone to transcriptional leakage that may be present in cells.

### Combining transcription and post-translation controls improves the robustness of the oscillator

We proposed to combine both a transcriptional and post-translational control for a more robust oscillator. The basic idea is that although the post-translation-based model, M2, has much worse oscillation in a stochastic condition (higher normalized decay rate) compared to M1, it will not have any leakage problem, since the transcription in M2 is not regulated. Instead, post-translational control may provide another layer of regulation for the repression leakage that occurred at the transcription level. To test this idea, we built a simple model, M5, that combined both M1 and M2 and performed a similar test as described above (Figure 1B). We found that 731 out of 1,000 parameter sets (73.1%) yielded sustained oscillation under a noisy condition, with a normalized decay rate median value of 1.68 (Figure 1C). Although the median value was still higher than M1, unlike M2, model M5 had a wider normalized decay rates distribution. It implied that M5 could still achieve robust oscillation under correct combination of parameters. Moreover, we also found that M5 was much more robust as compared with M1 under the basal leakage condition, such that 38.1% of parameter sets were still oscillating under 5% basal leakage (Figure 1E and Figure S2C). In addition, we also found that 16.3% of the parameter sets in model M5 were still able to oscillate even at 25% leakage level (compared to only 0.3% of parameter sets in model M1) (Figure S2A and Figure S2C). These results imply that model M5 may have a unique property to handle the basal leakage.

Adding a post-translational control in the transcriptional-based oscillator may help the system regulate the leakage in the transcriptional repression, such that it can still push the protein level back despite the leakage in the mRNA level. To demonstrate this idea, we showed that although we could still observe obvious shifting in the mRNA distribution for both models when we added a transcriptional leakage, the shifting of protein distribution in model M5 was kept at minimum compared to the shift observed in M1 (Figure 2). As a result, the X transcriptional inhibition of Z, for example, was also altered minimally in model M5 (Figure 2B, bottom). However, shifting the protein distribution of X greatly altered the X transcriptional inhibition of Z in M1 and locked it in the tight repressing state, which broke the oscillation (Figure 2A, bottom). Thus, our results suggested that combining both transcriptional and post-translational controls improve the robustness of the oscillator by controlling protein quantity when the mRNA transcribed is increased.

**Figure 2.**
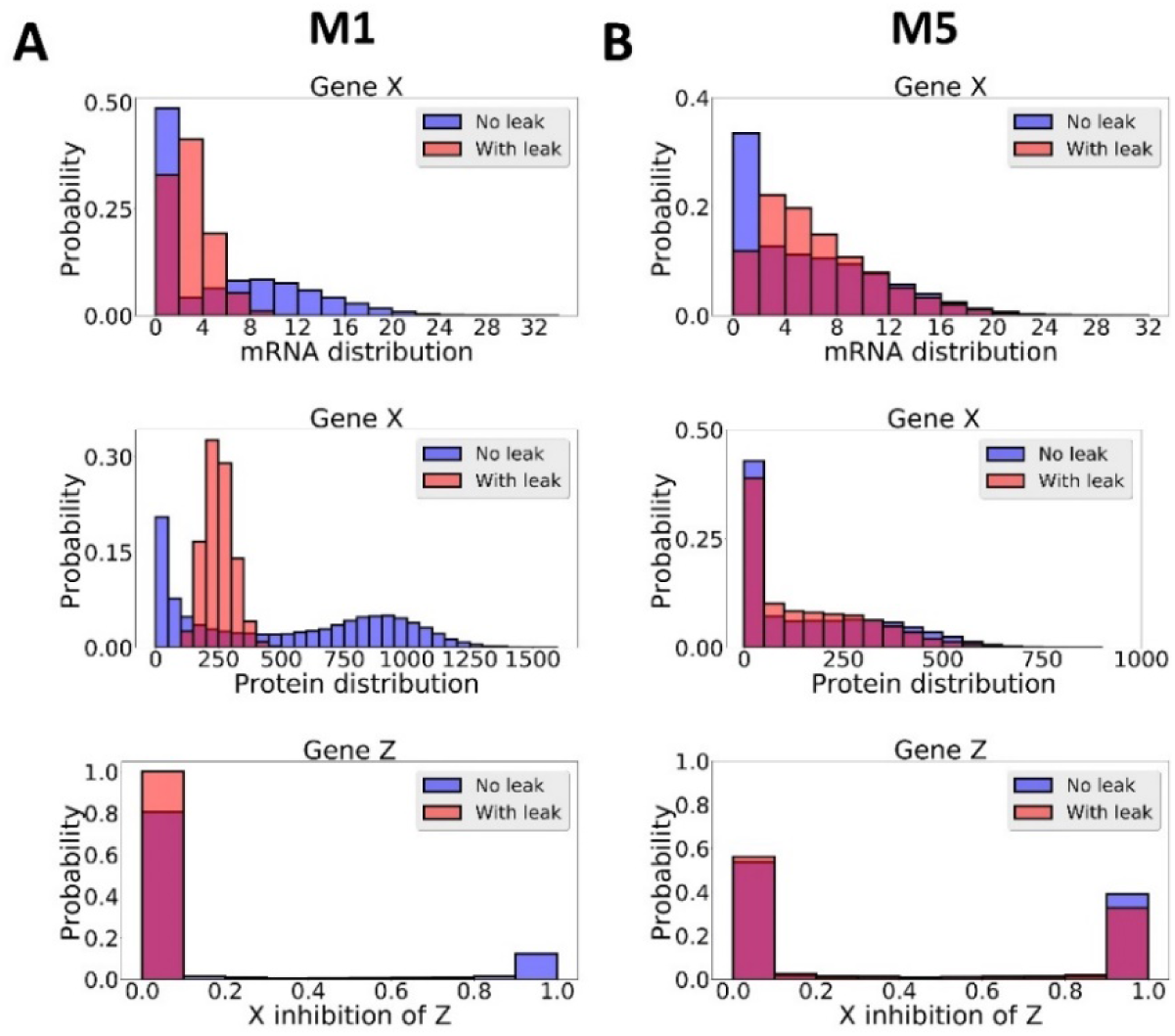
The distribution of mRNA (upper panel), protein (middle panel), and transcription activity (the Hill function) of X inhibition of Z (lower panel) for one randomly chosen parameter set in M1 (A) or M5 (B).

### Coupling degradation control to a repressilator using feed-forward loops gives better control to the leakage problem

To check the generality of our finding, we next expanded our model by taking into account other possible network structures and performed similar analyses as we did previously (Methods). For simplicity, but without loss of generality, we limited our analysis by keeping the repressilator as the core and targeted degradation (to be more specific 26S proteasome degradation) as the post-translation regulation. Other post-translational controls, such as phosphorylation, can also be modeled with similar mathematical forms to this degradation control, except for both the positive or negative effects to the system. With this formulation, we limited our analysis to four possible network structures from four different network motifs, which are type-3 coherent feed-forward loop (CFFL), type-2 incoherent feed-forward loop (IFFL), negative feedback (NF), and positive feedback (PF) (Figure 3A).

**Figure 3.**
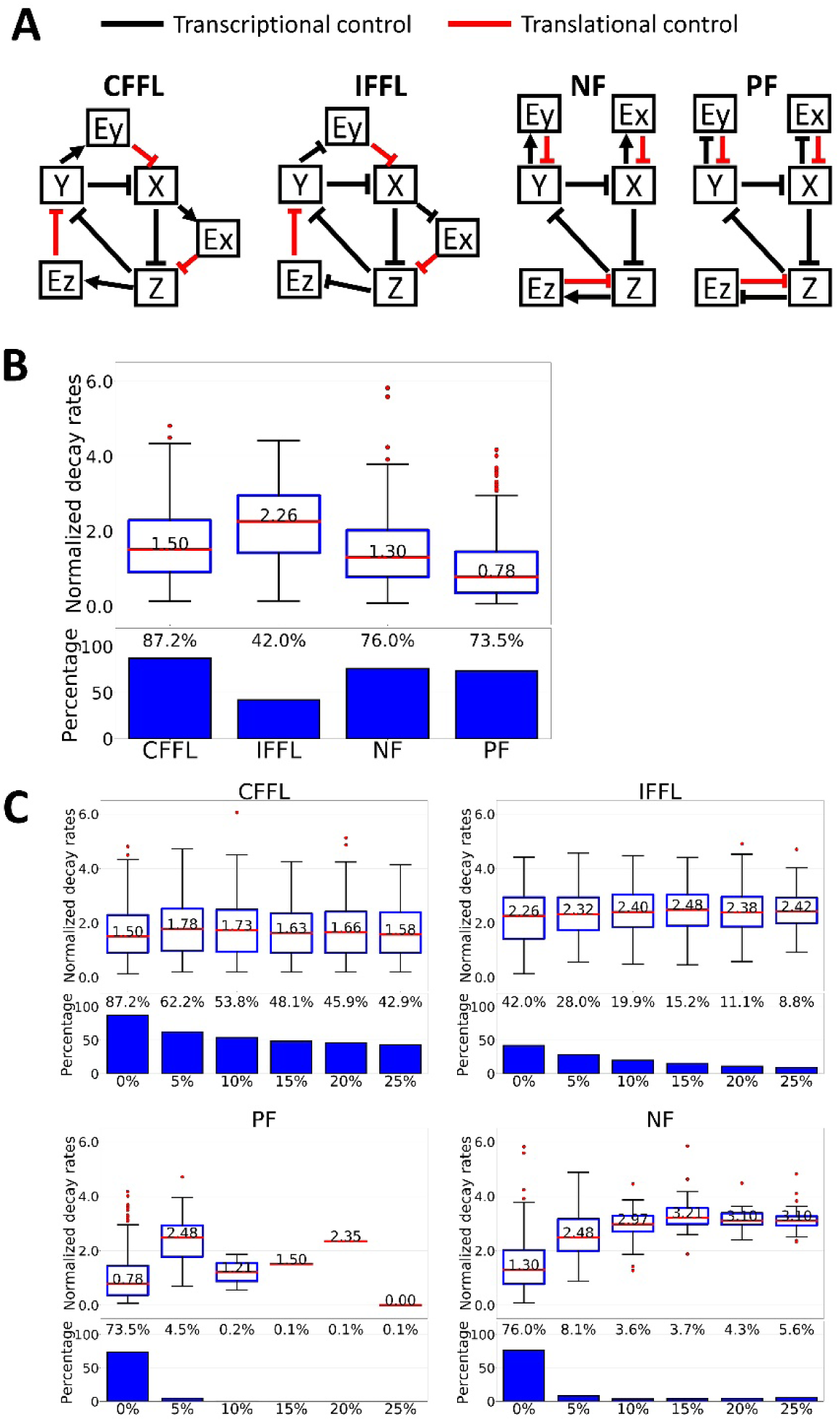
Robustness test of several network motifs in coupling the degradation control to a repressilator. (A) Schematic representation of the tested model. (B, C Upper panel) Box plot representing normalized decay rates of each tested model for 1000 randomly select parameter sets in the absence (B) or presence (C) of transcriptional leakage. Red lines indicate the median, and box edges indicate the 25th (Q1) and 75th (Q3) percentiles. Whiskers are plotted at 1.5*(Q3-Q1). (B, C Lower panel) Percentage of parameter sets that showed sustained oscillation under stochastic simulations.

In the absence of transcriptional leakage, we found that the number of parameter sets that could produce sustained oscillation under the noisy condition were still lower in all tested model compared to M1 (Figure 3B). Moreover, the median value of normalized decay rates in CFFL, IFFL and NF were still higher than M1, which is consistent with our previous observation using M5 (Figure 3C and Figure 3B). Of note, the PF model showed a better normalized decay rate compared to M1. This result is actually consistent with previous findings showing that coupling positive and negative feedback can create a more robust oscillation (61-64). However, coupled positive and negative feedbacks must occur at transcription and post-translation levels. Otherwise, a decreased robustness in the oscillation was observed (M3 in Figure 1C).

In the presence of transcriptional leakage, we again found a similar observation as in model M5 for CFFL and IFFL models, such that 62.2% and 28% of parameter sets were still oscillating under 5% basal leakage, respectively. However, both PF and NF models failed to survive with transcriptional leakage (Figure 3C). This finding is intriguing because both PF and NF models have similar degradation controls. To have better understanding of these results, we broke down the model into smaller network motifs and tested the effect of adding transcriptional leakage on both the input and output genes (Figure 4).

**Figure 4.**
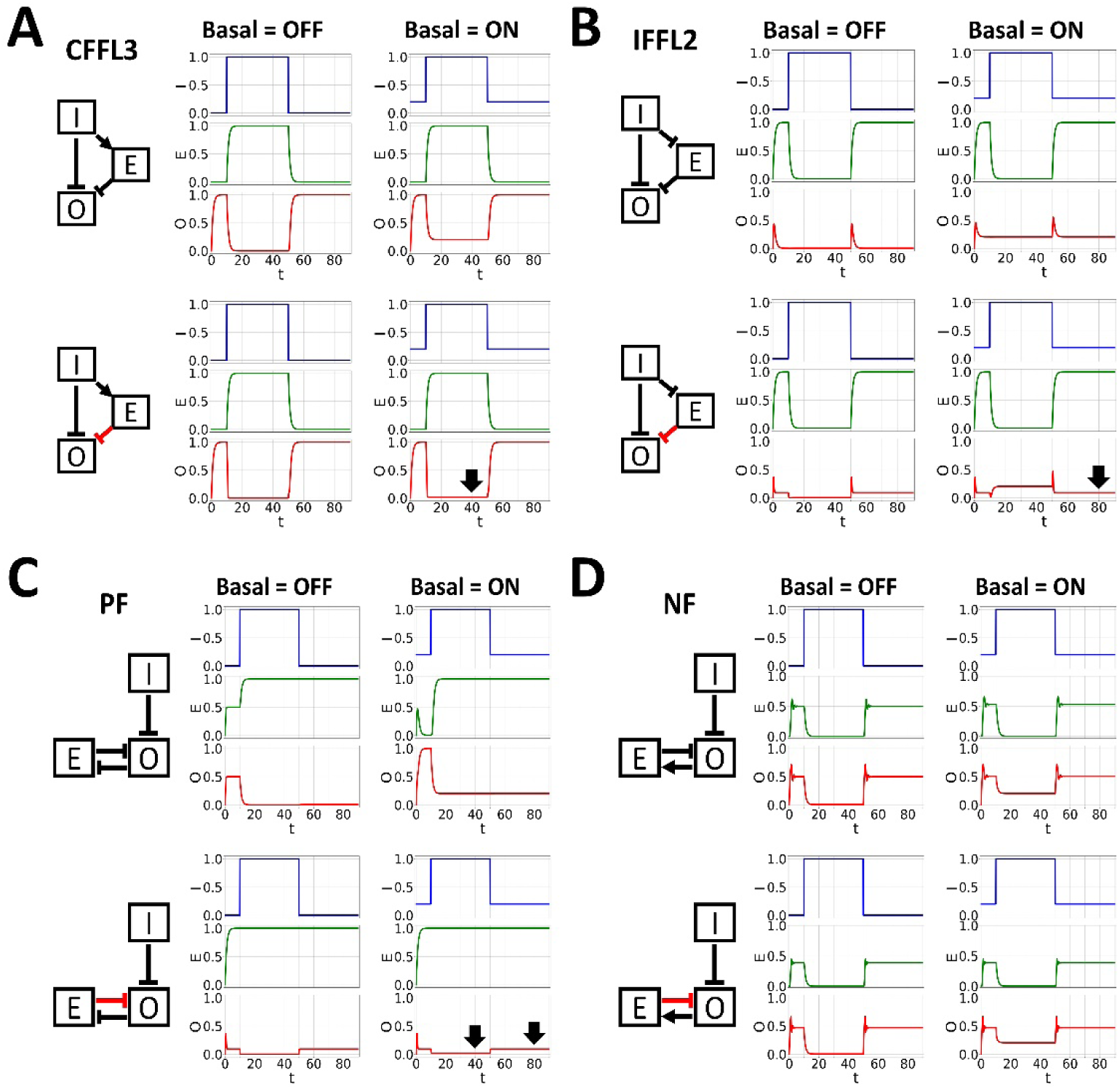
The dynamics of (A) type-3 coherent feed-forward loop, (B) type-2 incoherent feed-forward loop, (C) positive feedback, and (D) negative feedback in controlling transcriptional leakage. In upper panels, all genes are regulated through transcriptional controls. In lower panels, the E regulation of O was changed to a degradation control (indicates by red arrow). Black thick arrow highlights the filtering effect of a degradation control in the given network motif.

Here, we tested the added motif in assisting the oscillation by analyzing it locally. I and O denote input and output genes, respectively, which are part of the core elements in the repressilator (X, Y, or Z). The effect of translation control is considered advantageous if the turned-on basal expression in I leads to a similar status in O as if the basal were not there. As seen in Figure 4, the FFL motif is able to control the leakage when the input gene is ON, because the degradation control, E, is only ON when I is ON. Hence, it can push back the transcriptional leakage that occurs in the output gene (Figure 4A and Figure S3). In contrast, in the IFFL motif, gene E is expressed only when gene I is OFF. Thus, IFFL can only control the leakage in O when I is OFF (Figure 4B and Figure S4). For the PF motif, the leakage was actually controlled in both ON and OFF states of I (Figure 4C and Figure S5). However, PF motif has another problem, in which gene O is mostly kept in the OFF state (Figure S5 basal OFF panel). To have a higher amplitude of O, the PF motif requires the E regulation of O (K_Oe) to be weak (> 0.8, Figure S5). However, when the E regulation of O is weak, the ability of the degradation control to push back the basal leakage is also weaker. Hence, we see no filtering ability in our PF model (Figure 3C). For the NF motif, we also cannot find any filtering ability of the degradation control on the transcriptional leakage (Figure 4D and Figure S6). In this network, the degradation control E is only accumulated when the target gene O is ON. Since the transcriptional leakage occurs mostly when O is OFF, the degradation control has almost no effect, which we can see in the NF model result above (Figure 3C). Thus, our results suggest that in the presence of transcriptional leakage, coupling 26S proteasome degradation using feed-forward loops can help the repressilator to oscillate robustly.

### Our general models are able to reproduce the observed behavior in plant and mammalian clocks

From our observations above, we speculate that the 26S proteasome degradation could be important for the circadian clock to cope with transcriptional leakage and achieve robust oscillations in cells. To demonstrate this idea, we tried to replicate the dosage-dependent effect of the *TOC1* and *PRR5* degradation control, *ZEITLUPE* (*ZTL*), on the dynamics of the *Arabidopsis* clock (1). In 2004, Somers *et al.* showed that *ZTL* level controlled the amplitude and period of circadian oscillations (1). Moreover, it also showed that a different *ZTL* dosage affected the robustness of the oscillation, which could be seen from a higher relative amplitude error (RAE) value (Figure 5A). We hypothesized that the increase in RAE is due to the shifting of the protein level in degradation control, where the system failed to push back the basal leakage under the mutant condition but greatly reduced the amplitude of *TOC1* and *PRR5* proteins under high continuous overexpression.

**Figure 5.**
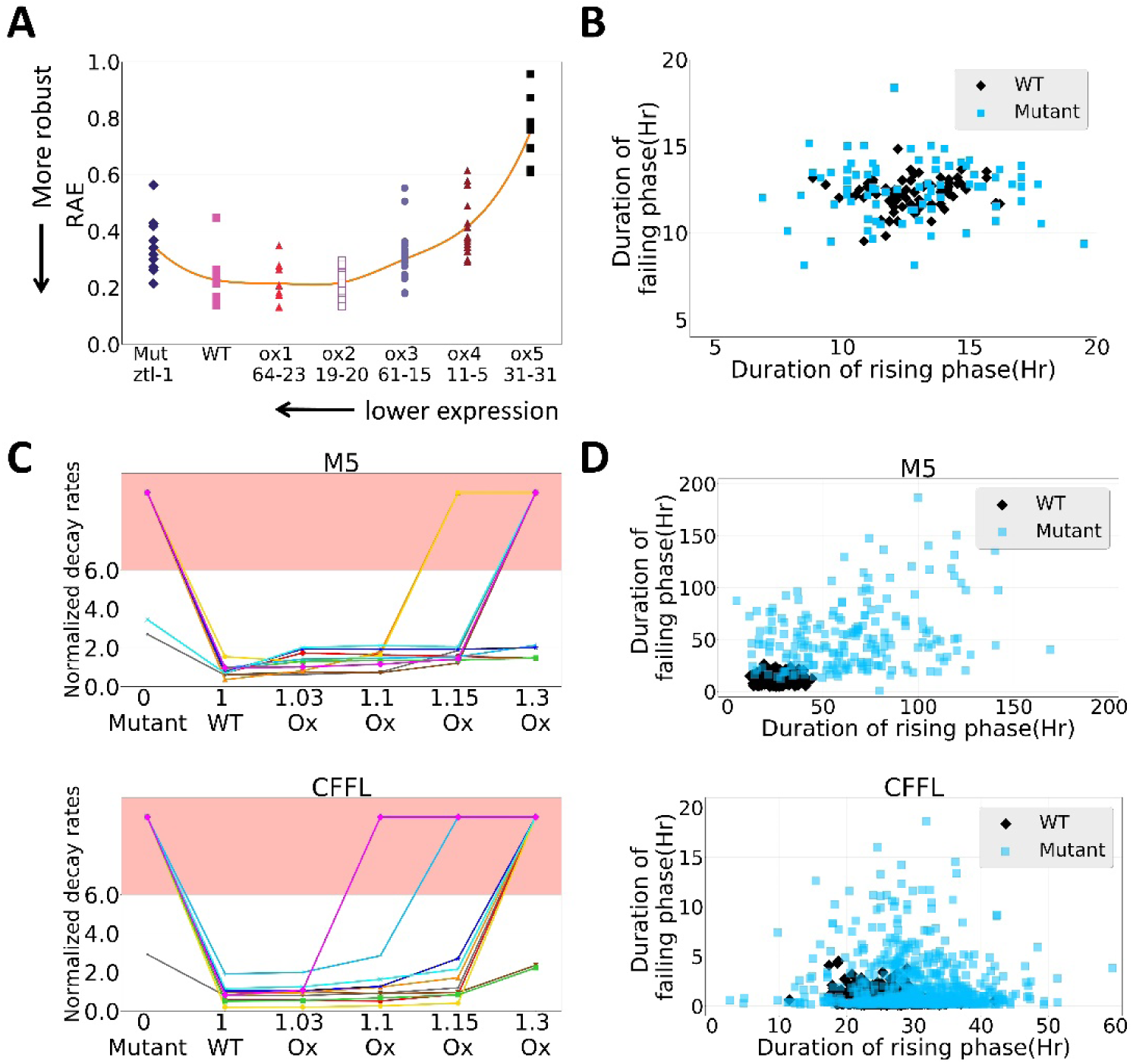
Our general models were able to replicate several observed experimental results. Experimental result of dosage-dependent effect of proteasome degradation control (*ZTL*) to the robustness of the oscillator. Data were obtained and redrawn from Somers *et al.* 2004 (1). (B) Experimental result of *β-Trcp2* mutant cells showed unstable rhythms and reduced amplitude of *PER* gene oscillations. Data were obtained and redrawn from D’Alessandro *et al.* 2017 (2). (C) The effect of changing the degradation control in model M5 (upper panel) and FFL (lower model) during our simulations. Each line represents one parameter set. The red part of the plot indicates the non-oscillating region. (D) The effect of mutating the degradation control to the rising and failing phases of model M5 (upper panel) and FFL (lower panel) during our simulations. The plot was drawn from one randomly selected parameter set.

To validate our hypothesis, we tried to replicate the dosage-dependent effect observed in the experimental using our general model M5. Here, we varied the X post-translational regulation of Z by mutating or constantly overexpressing it during simulations (Method). Under the mutant condition, the decay rates were generally increased (Figure 5C, upper panel) and the number of parameter sets that yielded oscillations were reduced from 28.1% to 5.4% under 10% leakage (Figure S7A). This result is similar to the experimental observation and was expected because the mutation of the X degradation control of Z will increase the basal expression of Z due to transcriptional leak. Hence, the system will suffer like the model M1 we described above (Figure 2).

However, the results of the overexpression condition are less straightforward than the mutant condition. We found that similar to the experimental result, the trend of decay rates was also increasing whereas the number of parameter sets that yielded oscillation also decreased along with the level of overexpression (Figure 5C and Figure S7A). Intuitively, one may expect that with more degradation control, we will see a more robust oscillation since the system can have better control on the transcriptional leakage. Hence, to have a better understanding of these results, we again tried to study the dynamics of the system by comparing the distribution of the overexpression mutant with the will-type condition (Methods). Here, we found that adding a constant amount of targeted degradation will have a larger effect on the amplitude of protein Z rather than keeping the basal level of Z in the very low level. This decrease in Z protein level further affects the Z degradation control of Y, which increases the Y protein level. After that, the increase in Y protein greatly reduced the expression of gene X (Figure S8), which prevented the system from having robust oscillation. Furthermore, to rule out a possible limitation of using a simple model (such as M5), we performed a similar analysis using a more elaborate model, FFL. Here, we again obtained a similar result such that the robustness of the oscillation was reduced when we mutated or overexpressed the Ex degradation control of Z (Figure 5C, lower panel and Figure S8).

Lastly, we also tried to replicate the effect of mutating the E3 ligase gene on the dynamics of the mammalian clock. In 2017, D’Alessandro *et al.* showed that mutation of the E3 ligase gene, *β-Trcp2*, will perturb the balance of *PERIOD* (*PER*) degradation, which makes the clock unstable (Figure 5B) (2). As we discussed above, we also observed unstable oscillation when we changed the degradation control in our models. We believe that our previous insight can be used to explain the observed experimental results in the mammalian clock. Hence, we again performed a similar analysis as we did previously and found that mutation of the degradation control will indeed alter the duration of the rising and failing phase of the oscillation (Figure 5B and D). These results indicate that our hypothesis can also be found in a real oscillating system such as circadian clocks.

## Discussion

### Coupling E3 ligase to a negative feedback loop using feed-forward loop is commonly seen in the *Arabidopsis* circadian clock

Our results showed that combining two types of regulation using feed-forward loop can create a robust oscillator. In the *Arabidopsis* clock, several E3 ligases and their respective targets have been successfully identified. The first identified E3 ligase was an F-box protein, *ZEITUPLE (ZTL). ZTL* was reported to be involved in target degradation of both *PRR5* and *TOC1* proteins (31, 39). Although *ZTL* mRNA is consecutively expressed, its protein still oscillated with threefold change in amplitude (65). This oscillation may be mediated through a protein–protein interaction of *ZTL* with *GIGANTEA (GI*) protein (66). Also, *CCA1* and *LHY* can bind and repress the expression of *GI.* Hence, together with ZTL, they form a feed-forward network motif (Figure 6A).

**Figure 6.**
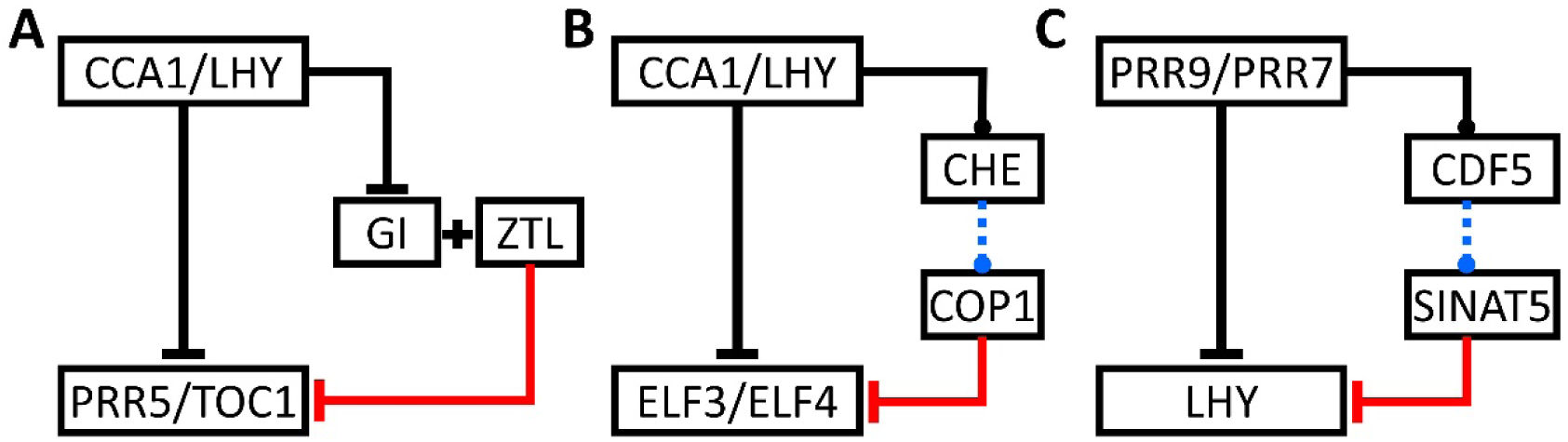
Currently known (A) or predicted (B and C) regulators of E3 ligase in the plant’s clock form a feed-forward network motif. Black and blue lines represent transcriptional regulations, and the red line represents the degradation control. Solid lines indicate that the data were derived from experimental results, whereas dashed lines indicate that the data were derived from TF binding prediction tools, PlantTFDB (67) and PlantPAN3 (68).

Next, a RING-type E3 ligase protein, *constitutive photomorphogenesis 1 (COP1)*, has also been reported to degrade other important clock genes, *ELF3* and *ELF4* (41, 69). *ELF3, ELF4* and *LUX* have been reported to form a protein complex, called *evening complex (EC)*, and the loss-of-function mutant of any of these three proteins will lead to arrhythmic behavior (28, 70-72). Previously, *CCA1* and *LHY* were reported to bind and repress the expression of both *ELF4* and *ELF3* (17, 73, 74). However, little is known about the regulator of *COP1* proteins. From our results, we hypothesized that the clock system could have the advantages that we mentioned above if *CCA1* and *LHY* can directly (or indirectly) regulate COP1. Hence, we performed a quick analysis combining ChIP-seq data for *CCA1* (17, 73) and *LHY* (74) with TF binding prediction tools, PlantTFDB (67) and PlantPAN3 (68), to predict the direct or indirect regulation of *CCA1*/*LHY* to *COP1*. Interestingly, our analysis suggests that a well-studied transcriptional factor, *TCP21 (TCP21/CHE)*, could bind to the *COP1* promoter region and both *CCA1* and *LHY* could also bind to the *TCP21* promoter region. Together, they form an indirect feed-forward network motif (Figure 6B).

Finally, another RING-type E3 ligase protein, *SINAT5* was also reported to interact and degrade *LHY* protein in the plant (36). However, similar to *COP1*, little is known regarding the regulator of *SINAT5*. Hence, we performed a similar analysis as we did for *COP1*, only now we used ChIP-seq data for *PRR9* and *PRR7* (75). We found that *cyclic dof factor5 (CDF5)* could bind to the SINAT5 promoter region, whereas both *PRR9* and *PRR7* were found in the *CDF5* promoter region. Together, they form another indirect feed-forward network motif (Figure 6C).

Although these predictions are yet to be verified experimentally, they provide other evidence that strengthens our simulation results. Previously, several mathematical models have shown that the core oscillator of the *Arabidopsis* clock consists of four groups of genes, the early morning phase genes (*CCA1/LHY*), the daytime/noon phase genes (*PRR9/PRR7*), the afternoon/dusk phase genes (*PRR5/TOC1*), and the nighttime/midnight phase genes (*EC*) (24-26, 48). Among these four groups, three may be regulated by coupled repression and degradation through feed-forward loops (Figure 6). To the best of our knowledge, the corresponding E3 ligases for the noon phase genes, *PRR9* and *PRR7*, are still elusive. However, several studies have shown that both genes are still regulated by the 26s proteasome degradation pathway (38, 76). It will be interesting to see whether these two genes are also under similar regulation. Thus, these findings imply that coupling E3 ligase to a negative feedback using a feed-forward loop is probably common in the *Arabidopsis* clock.

### Transcriptional leakage is commonly seen in cells

Transcription leakage (or basal transcription) commonly occurs in cells. For example, several systems like CpxR in *Escherichia coli* and VirG in *Agrobacterium tumefaciens* have been reported to have high basal expression (77-79). Moreover, Yanai *et al.* also showed that the expression of many tissue-specific genes can “overflow” into neighboring genes that have no function in the respective tissue (59). This observation implies that leakiness of the promoters may be evolutionarily neutral (60).

In this study, we showed that leakage can be a serious issue in oscillating systems like the circadian clock. In cells, this problem is probably handled by having a dual transcription and post-translation control, which gives another layer of regulation for cells to deal with the leakage problem. Actually, in line with our findings, many synthetic systems have emphasized the importance of combining both transcription and post-transcription/translation controls in creating a robust oscillator (78, 80-83). For instance, Tigges *et al.* showed that a synthetic mammalian oscillator based on transcription (sense) and a post-transcription control (antisense) can create an autonomous, self-sustained and tunable oscillator (81). In another study, Danino *et al.* showed that coupling transcription (LuxR) and degradation control (AiiA) with global intercellular coupling (AHL) generates synchronized, robust oscillation (82). These observations strengthen our finding that coupling transcriptional and degradation control may be important in controlling transcriptional leakage in cells.

However, although leaky expression is often considered as unwanted noise in cells, it can also be useful in some systems. For instance, in the competence development of *Bacillus subtilis*, basal expression of the master regulator (*comK*) must pass a critical threshold in order to trigger its autoactivation, which will lead to a bistable pattern in *B.subtilis* cells. This bistability will further create heterogeneity in cell populations, which can benefit the population by providing better-adapted phenotypic variants for a given perturbation (84). In contrast with this finding, Ingolia and Murray in 2007 showed a different role of basal expression in creating bistability. Using a budding-yeast pheromone response system, they could make this system bistable by reducing the basal expression (85). Hence, depending on the evolutionary process, basal expression can be considered an important aspect or a noise in different biological systems.

## Material and Methods

### Model Representation

All models used in this study are described by a set of ordinary differential equations (ODEs) for the simulation under continuous light. In general, each gene was represented as:

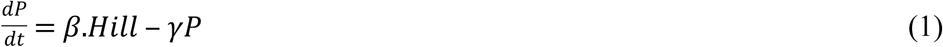

where P represents the dimensionless concentration, which can be any genes depending on the model. Here, β represents the total production rate, whereas γ is the total degradation rate. The nondimensionalization process involved choosing a constant value for each component, denoted as P_0_. P_0_ was defined as the maximum steady state of gene P, which is the ratio of total production rates over total degradation rates. Hill represents the Hill function, which describes the effects of upstream regulation as

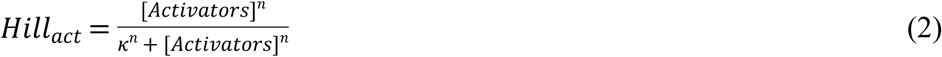

for the activating process and

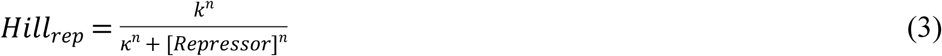

for the repression process. As mentioned previously, the Hill input function is a monotonic, S-shaped function, which is used to describe the effect of the transcription factor on the transcription rate of its target gene (86). Here, κ represents the concentration of the activator or repressor needed to achieve half maximal effect. It is related to the binding affinity between the transcriptional factor gene and its site on the promoter region (86). The *n* represents the Hill coefficient that governs the steepness of the input function, which is related to sensitivity of the process in the cell. Employing *n* allows us to describe ultra-sensitivity of many cellular processes, such as multisite phosphorylation, stoichiometric inhibitor, cooperativity, reciprocal regulation, and substrate competition (87). Previously, several clock proteins have been shown to form a dimer (29, 88, 89). Because of this reason, many mathematical models set *n* = 2, which corresponds to this dimerization process (24, 48, 90). However, in this study, we allowed *n* to be > 2 to accommodate other possible regulations (Table 1). Following previous studies, we used an ‘AND’ gate to describe a combination of two or more source of regulations, where the two Hill functions are multiplied (24, 26, 48).

**Table 1.**
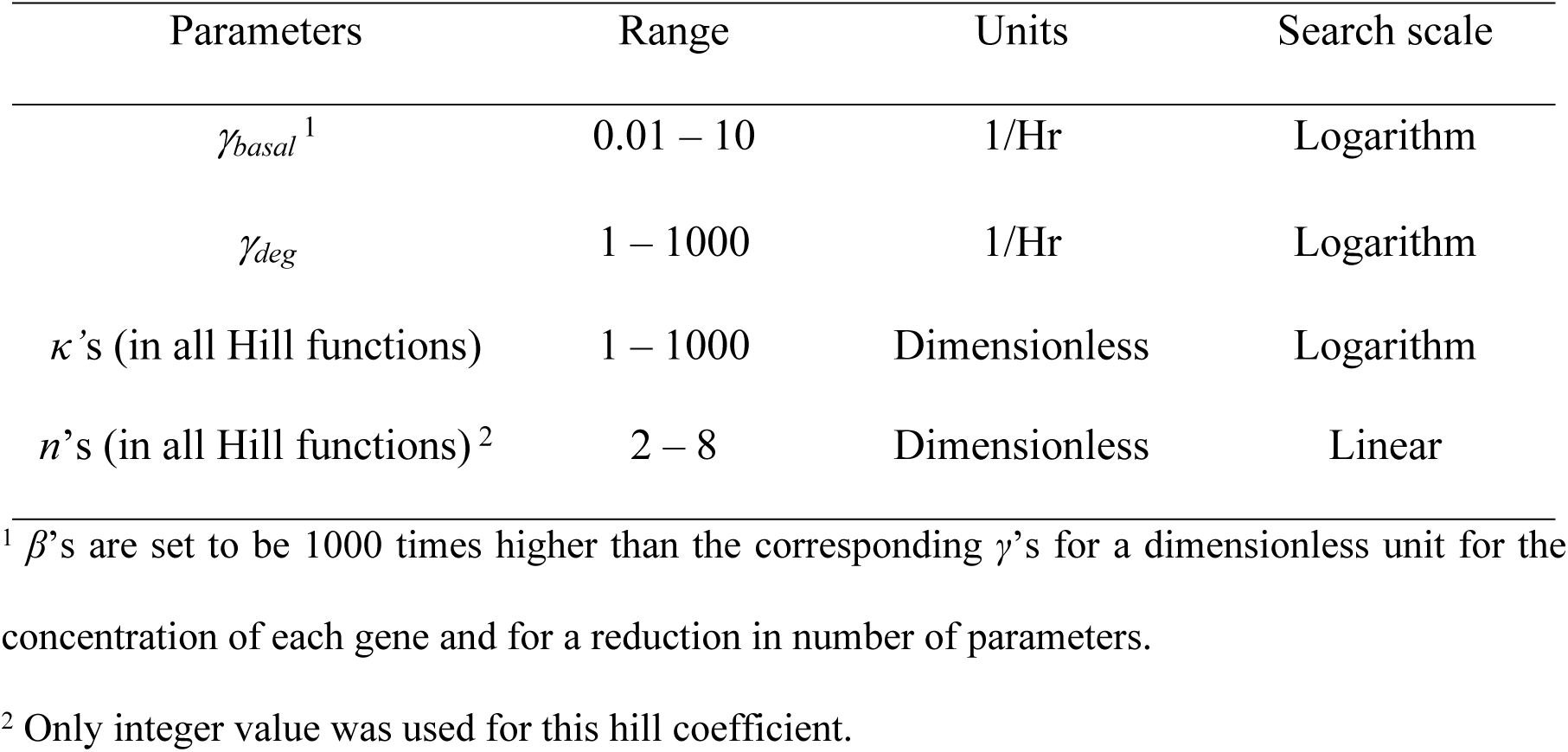
Search ranges for parameters.

| Parameters | Range | Units | Search scale |
| --- | --- | --- | --- |
| $\gamma_{basal}$<sup>1</sup> | 0.01 – 10 | 1/Hr | Logarithm |
| $\gamma_{deg}$ | 1 – 1000 | 1/Hr | Logarithm |
| $\kappa$'s (in all Hill functions) | 1 – 1000 | Dimensionless | Logarithm |
| $n$'s (in all Hill functions)<sup>2</sup> | 2 – 8 | Dimensionless | Linear |

<sup>1</sup> $\beta$ 's are set to be 1000 times higher than the corresponding $\gamma$ 's for a dimensionless unit for the concentration of each gene and for a reduction in number of parameters.
<sup>2</sup> Only integer value was used for this hill coefficient.

Finally, we also fixed the maximum possible steady-state concentration of each component to 1000 molecules per cell by assuming the Hill function as its maximum possible value, 1. In this way, the maximum production rates (*β*’s) were fixed to be 1000 times higher than the total degradation rates (*γ’*s). The time *t* in the current work was in the unit of hours. After obtaining a regularly oscillating parameter set in the wild-type setting, we re-scaled all the time-related parameters such that the oscillation period is 24 hr.

- **Model M1: transcriptional control based repressilator**

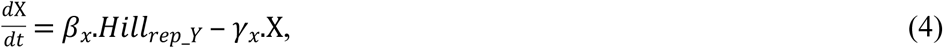

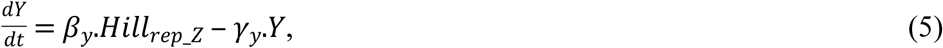

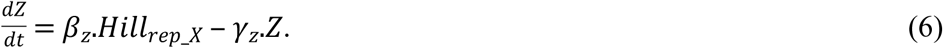
- **Model M2: Post-translational control based repressilator**

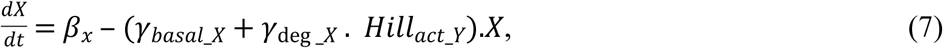

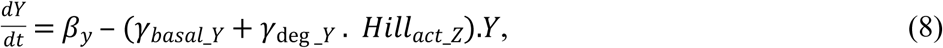

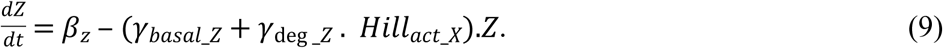 We assumed that the target degradation happened much faster than basal degradation. Hence, the searching space for γ_deg_ was set 2 orders higher (Table 1).
- **Model M3: Transcriptional control based repressilator with additional positive feedback loop** In model M3, equation (4) was modified into equation (10),

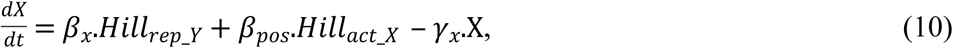

whereas the other two equations remained the same (equations (5) and (6)). There were two additional parameters (the corresponding production rates (β_pos_) and *κ* value in the new Hill function *Hill*_*act_X*_) added in this model. The parameters were treated similarly as those in M1.
- **Model M4: Post-translational control based repressilator with additional positive feedback loop** For model M4, equation (7) was modified into equation (11),

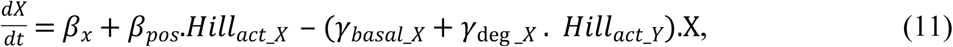

while other two equations (equations (8) and (9)) were kept the same. There were again two additional parameters added in this model. The parameters were treated similarly as those in M2.
- **Model M5: coupled transcriptional and post-translational control-based oscillator** For model M5, we combined the regulation of model M2 into model M1 and described it as:

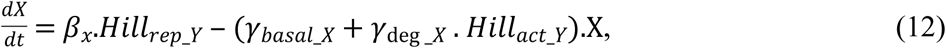

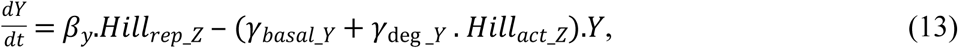

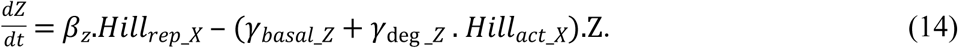 Although the transcriptional and post-translational regulations are usually controlled by different genes, for simplicity, we assumed that it might be originated from the same transcriptional regulator. For instance, to post-translationally modify protein Z, gene X might actually need to activate another protein, X’, that can bind to and modify protein Z. However, if we assume that the expression of gene X’ is solely dependent on gene X and this activation process happens fast enough, we can omit gene X’ in our model and replace this regulation by a hill function. By doing so, we can greatly reduce the complexity of the model M5.
- **Various variant of coupled transcription and post-translation model** To overcome the limitations of using simplified models, we next tried to elaborate our model by taking into consideration several types of regulations that have been commonly discussed in the other studies. Here, we tested four different network motifs, Coherent feed-forward loop (FFL), Incoherent feed-forward loop (IFFL), negative feedback loop (NF), and positive feedback loop (PF).
  1. ***Feed-Forward Loop (FFL)***

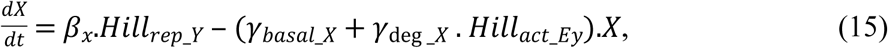

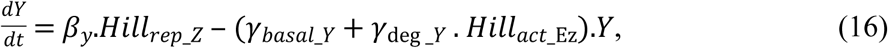

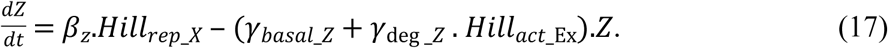

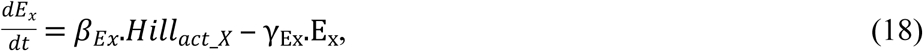

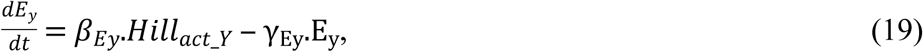

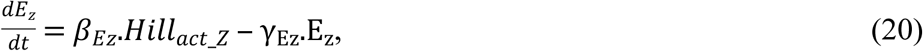
  2. ***Incoherent Feed-Forward Loop (IFFL)***

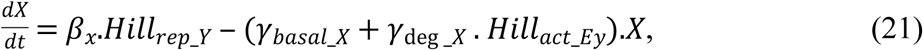

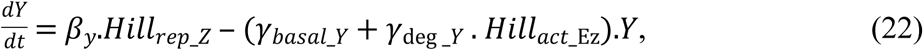

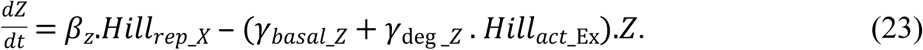

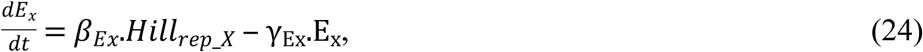

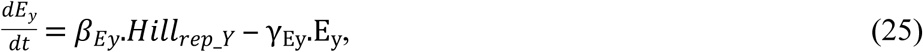

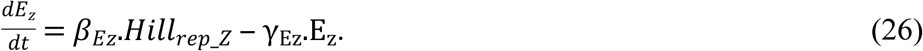
  3. ***Negative Feedback (NF)***

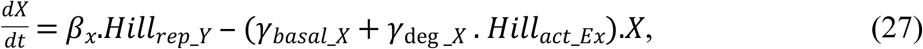

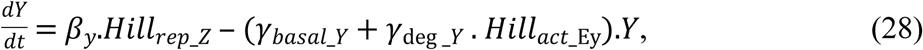

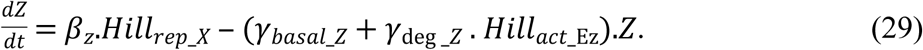

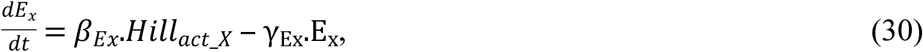

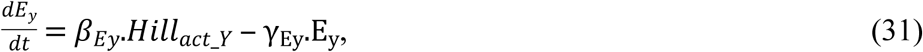

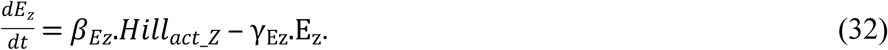
  4. ***Positive Feedback (PF)***

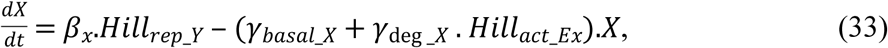

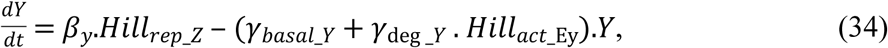

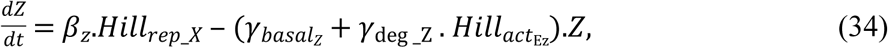

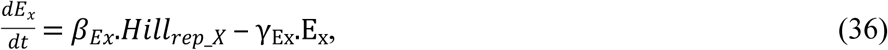

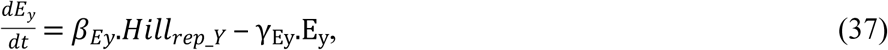

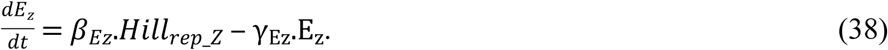

### Searching, propagation, and selection process

All independent parameters in each model were obtained by random searches, propagated, and screened for regular oscillation. The search was performed at a logarithmic scale across three orders of magnitude, for γ’s and κ’s, and a linear scale for *n* (Table 1). Here, a parameter set is selected if the trajectory can oscillate regularly, defined by examining the period and amplitude change in each cycle. We calculated the relative difference in period and amplitude change for each cycle, defined as 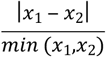, where *x*_1_ and *x*_2_ are the period or amplitude calculated from two consecutive cycles. An acceptable regular oscillation was defined as that with less than 5% relative change for more than 10 cycles. For all searching, we used similar initial conditions, which is 10% of maximum possible steady-state concentration. Finally, all simulations were performed by using Python 3.6 (Anaconda 4.4.0).

### Stochastic simulations

For all models, the stochastic simulation involved the Gillespie algorithm (53, 91). Here, each gene is described in three different levels: gene, mRNA, and protein levels. For instance, gene X in model M5 (equation (12)) was described as:

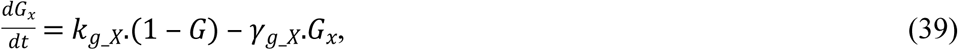

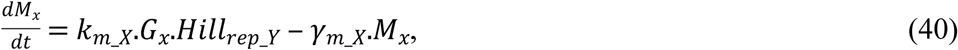

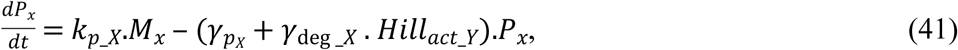

where *G* is the fraction of active gene for transcription, *M* is the amount of mRNA and *P* is for proteins. *k* represents the activation or production rates and γ is the deactivation or degradation rate. It has been shown previously that both mRNAs and protein are produced in bursts (92, 93). Following previous study, we defined the ratio between 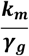 as the mean burst frequency (B_m_) and the ratio of 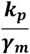 as mean burst size (B_p_) (94). For simplicity, but without loss of generality, we fixed our B_m_ and B_p_ into 2 and 10, respectively. Last, we also set γ_g_ and γ_m_ at 100 and 10 times of γ_p,,_ which satisfies the condition of burst production in both mRNA and protein levels as described previously (94). By doing so, we can obtain all necessary propensity functions from our ODE equation without any additional parameter value to be searched (Table 2).

**Table 2.** Summary of parameter conversion (using gene X of model M5 as an example)

| Propensity function | Corresponding values |
| --- | --- |
| $\gamma_{p\_x}$ | $\gamma_x^1$ |
| $\gamma_{m\_x}$ | $\gamma_{p\_x} * 10$ |
| $\gamma_{g\_x}$ | $\gamma_{p\_x} * 100$ |
| $k_{g\_x}$ | $\frac{\beta_x^1}{Bm * Bp}$ |
| $k_{m\_x}$ | $Bm * (k_{g\_x} + \gamma_{g\_x})$ |
| $k_{p\_x}$ | $Bp * \gamma_{m\_x}$ |
| $Hill_{rep\_y}$ | $Hill_{rep\_y}^1$ |
| $\gamma_{deg\_x}$ | $\gamma_{deg\_x}^1$ |
| $Hill_{act\_y}$ | $Hill_{act\_y}^1$ |

<sup>1</sup> Parameter value obtained from ODEs simulation.

### Measuring the oscillations under stochastic simulations

There are several ways to measure the “goodness” of oscillatory behavior (95, 96). However, in this study, we make use of the autocorrelation function to measure the “goodness” of the oscillations. Let P_(M.Δt)_ be a time series data, corresponding to one protein component in our simulations, with length of M times Δt (in our simulation, Δt is fixed into Hr). Mathematically, the autocorrelation function of P_(M.Δt)_ is defined as:

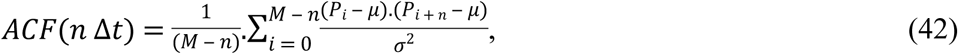

where *n* can be any value between 0 to M-1. In our simulation, we fixed our *n* such that it is between 0 and 2400 (equivalent to 10 days). Using this equation, the maximum value of equation (42) is when *ACF*(0), which is 1. If P_(M.Δt)_ is the output of deterministic simulation with sustained oscillation, then equation (42) will also oscillate in a sustained manner. This kind of oscillation can be estimated by using equation (43),

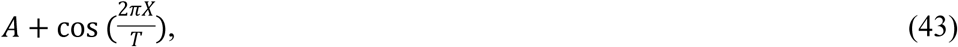

where *X* is the autocorrelation function value, and *T* is the period of the oscillation. However, if P_(M.Δt)_ describes a realization of a stochastic oscillatory process, equation (42) will show a damped oscillation. In this case, this dampened oscillation can be estimated by equation (44),

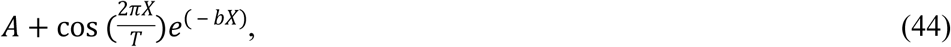

where *b* represents the damping rate or characteristic time of the decay of the autocorrelation function (96). Notice that parameter b will still carry the time dimension. Hence, it can be nondimensionalized by dividing it with the period parameter, T. In this study, we can even further simplify this estimation process by just fitting the dampened autocorrelation function with an exponential function,

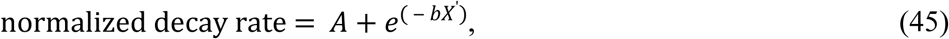

where X’ represents all the peak value obtained in the autocorrelation function.

### Modified stochastic simulations

- **Adding transcriptional leakage** To add a transcriptional leakage in our stochastic simulations, we modified equation (40),

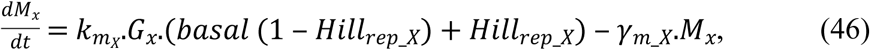

and applied it to all genes in each model. Other equations (equations (39) and (41)) were kept the same. We argue that this representation is more realistic than simply adding a constant basal expression in the mRNA level, because with this modification, the maximum value of the hill function is kept.
- **Mutation and overexpression of degradation control in M5** For the overexpression test of model M5, we modified the X degradation control of gene Z (*Hill*_*act_X*_ in equation (14)) from equation (2) into equation (47),

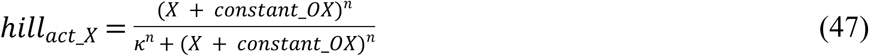

with other equations kept the same. In this analysis, we cannot simply overexpress gene X, because it would also affect the gene X transcriptional regulation of gene Z (*hill*_*rep_X*_ in equation (14)). Next, we varied the value of *constant_OX*, from 3% to 30%, based on the maximum possible steady state of ODE oscillation (which is 1000). For the null-mutant test, we simply set *γ*_deg *_Z*_ in equation (14) to 0.
- **Mutation and overexpression of degradation control in other models** For the other models (FFL, IFFL, NF, PF), the genetic perturbation tests were done by scaling up (for overexpression) or scaling down (for null-mutant) the production rates (*β*) of gene Ex (for FFL and IFFL) or gene Ez (for NF and PF), while keeping the same degradation rates (*γ*). We again varied it from 3% to 30%, based on the maximum possible steady state of ODE oscillation (which is 1000).

### Testing the effect of different network motifs in the coupling degradation control to transcriptional oscillator

To test whether a certain network motif has better capability in controlling basal expression, we built partial models consisting of only the type-3 coherent feed-forward loop (FFL), type-2 incoherent feed-forward loop (IFFL), negative feedback (NF), and positive feedback (PF) in either ‘pure’ transcriptional or combined transcriptional and degradation controls.

Here, we measured the concentration of the output gene (O) in the presence (I:ON, t=40) or absence (I:OFF, t=80) of an input signal (I), which we called the basal OFF condition. For the basal ON condition, we added the leakage term on both input and output genes (leakage level = 0.2, or 20% of the maximum steady state) and re-measured the concentration of the output genes. After that, we calculated the normalized difference of output gene expression, which is defined as:

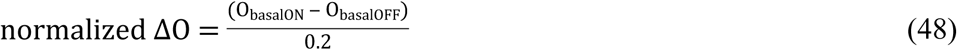

Finally, we subtracted the normalized ΔO of transcriptional model to the normalized ΔO of the combined transcription and post-translation model (Δnormalized ΔO) and report it as an indicator of how good a given motif is in controlling transcriptional leakage (Figure 4 and Figure S3-S6).

## Data Availability

All data generated or analyzed during this study are included in this published article (and its Supplementary Information files)

## Acknowledgments

We acknowledge support from Academia Sinica through the Investigator Award (AS-IA-106-M01), the Ministry of Sciences and Technology of Taiwan (grant no. MOST 105-2113-M-001 -009 -MY4), and a joint research grant from National Taiwan University and Academia Sinica for support of this research work. This work also benefited from events organized by the National Center for Theoretical Sciences.

## Author contributions

I.J., S.H.W., and C.P.H. designed the research, analyzed the data. I.J. and C.P.J. wrote the article. I.J and C.C.S.Y. developed the simulation method and performed the simulation. J.W.C., S.H.W., and C.P.H supervised the research project. All authors reviewed the manuscript.

## Supporting information captions

**Figure S1 Sample trajectories of mRNA (left), protein (middle), and auto-correlation function (right) of one randomly chosen parameter set in model M1 (a) and M2 (b).**

**Figure S2 The effect of transcriptional leakage on the robustness of an oscillator.** (Upper panels) box plot representing the normalized decay rates for each leakage level of model M1 (A), M3 (B), and M5 (C). (Lower panels) the percentage of parameter sets showing sustained oscillation under stochastic simulation. Red line indicates the median, and box edges indicate the 25th (Q1) and 75th (Q3) percentiles. The whiskers are defined as 1.5*(Q3-Q1).

**Figure S3 Contour plot representing the dynamics of type-3 coherent feed-forward loop in controlling transcriptional leakage under different parameter combinations.** I, E, and O representing the input, intermediate, and output genes, respectively. For simplicity, but without loss of generality, the production and the degradation rates of all genes was fixed at 1, the threshold value of I activation of E (K_Ei) at 0.5, and the Hill coefficient at 8. Furthermore, the leakage level (when Basal = ON) was fixed at 0.2 (or 20% of the maximum possible steady state).

**Figure S4 Contour plots representing the dynamics of type-2 incoherent feed-forward loop in controlling transcriptional leakage under different parameter combinations.** I, E, and O represent the input, intermediate, and output genes, respectively. For simplicity, but without loss of generality, the production and the degradation rates of all genes was fixed at 1, the threshold value of I inhibition of E (K_Ei) at 0.5, and the Hill coefficient at 8. Furthermore, the leakage level (when Basal = ON) was fixed at 0.2 (or 20% of the maximum possible steady state).

**Figure S5 Contour plots representing the dynamics of positive feedback in controlling transcriptional leakage under different parameter combinations.** I, E, and O represent the input, intermediate, and output genes, respectively. For simplicity, but without loss of generality, the production and the degradation rates of all genes was fixed at 1, the threshold value of O inhibition of E (K_Eo) at 0.5, and the Hill coefficient at 8. Furthermore, the leakage level (when Basal = ON) was fixed at 0.2 (or 20% of the maximum possible steady state).

**Figure S6 Contour plots representing the dynamics of negative feedback in controlling transcriptional leakage under different parameter combinations.** I, E, and O represent the input, intermediate, and output genes, respectively. For simplicity, but without loss of generality, the production and the degradation rates of all genes was fixed at 1, the threshold value of O activation of E (K_Eo) at 0.5, and the Hill coefficient at 8. Furthermore, the leakage level (when Basal = ON) was fixed at 0.2 (or 20% of the maximum possible steady state).

**Figure S7 Our general models showed similar dosage-dependent effect of proteasome degradation control on the robustness of the oscillator.** (Upper panel) box plot representing the normalized decay rates for each mutant condition of model M5 (A), and FFL (B). Red line indicates the median, and box edges indicate the 25th (Q1) and 75th (Q3) percentiles. The whiskers are defined as 1.5*(Q3-Q1). (Lower panel) The percentage of parameter sets showing sustained oscillation under stochastic simulation.

**Figure S8 The distribution of mRNA (first panel), protein (second panel), Hill function of transcriptional control (third panel), and Hill function of degradation control (last panel) for one randomly chosen parameter set in M5.** The distribution was collected from the wild type (WT) (blue) or over-expression condition (red) with 10% basal leakage.

## References

1. Somers DE, Kim W-Y, Geng R. The F-Box Protein ZEITLUPE Confers Dosage-Dependent Control on the Circadian Clock, Photomorphogenesis, and Flowering Time. The Plant Cell. 2004;16(3):769–82.

2. D’Alessandro M, Beesley S, Kim JK, Jones Z, Chen R, Wi J, et al. Stability of Wake-Sleep Cycles Requires Robust Degradation of the PERIOD Protein. Current Biology. 2017;27(22):3454–67.e8.

3. Cohen SE, Golden SS. Circadian Rhythms in Cyanobacteria. Microbiology and molecular biology reviews : MMBR. 2015;79(4):373–85.

4. Heintzen C, Liu Y. The Neurospora crassa Circadian Clock. Advances in genetics. Volume 58: Academic Press; 2007. p. 25–66.

5. Hardin PE. Chapter 5 - Molecular Genetic Analysis of Circadian Timekeeping in Drosophila. In: Stuart B, editor. Advances in genetics. Volume 74: Academic Press; 2011. p. 141–73.

6. Lowrey PL, Takahashi JS. Genetics of circadian rhythms in Mammalian model organisms. Advances in genetics. 2011;74:175–230.

7. McClung CR. CIRCADIAN RHYTHMS IN PLANTS. Annual review of plant physiology and plant molecular biology. 2001;52:139–62.

8. Kovac J, Husse J, Oster H. A time to fast, a time to feast: the crosstalk between metabolism and the circadian clock. Molecules and cells. 2009;28(2):75–80.

9. Gerstner JR, Yin JCP. Circadian rhythms and memory formation. Nat Rev Neurosci. 2010;11(8):577–88.

10. Dodd AN, Salathia N, Hall A, Kevei E, Toth R, Nagy F, et al. Plant circadian clocks increase photosynthesis, growth, survival, and competitive advantage. Science. 2005;309(5734):630–3.

11. Fu L, Kettner NM. Chapter Nine - The Circadian Clock in Cancer Development and Therapy. In: Martha UG, editor. Progress in Molecular Biology and Translational Science. Volume 119:Academic Press; 2013. p. 221–82.

12. Dupuis J, Langenberg C, Prokopenko I, Saxena R, Soranzo N, Jackson AU, et al. New genetic loci implicated in fasting glucose homeostasis and their impact on type 2 diabetes risk. Nat Genet. 2010;42(2):105–16.

13. Fonseca Costa SS, Ripperger JA. Impact of the Circadian Clock on the Aging Process. Frontiers in Neurology. 2015;6(43).

14. Nohales MA, Kay SA. Molecular mechanisms at the core of the plant circadian oscillator. Nat Struct Mol Biol. 2016;23(12):1061–9.

15. Alabadi D, Oyama T, Yanovsky MJ, Harmon FG, Mas P, Kay SA. Reciprocal regulation between TOC1 and LHY/CCA1 within the Arabidopsis circadian clock. Science. 2001;293(5531):880–3.

16. Adams S, Manfield I, Stockley P, Carré IA. Revised Morning Loops of the <italic>Arabidopsis</italic> Circadian Clock Based on Analyses of Direct Regulatory Interactions. PLoS ONE. 2015;10(12):e0143943.

17. Kamioka M, Takao S, Suzuki T, Taki K, Higashiyama T, Kinoshita T, et al. Direct Repression of Evening Genes by CIRCADIAN CLOCK-ASSOCIATED1 in the Arabidopsis Circadian Clock. Plant Cell. 2016;28(3):696–711.

18. Gendron JM, Pruneda-Paz JL, Doherty CJ, Gross AM, Kang SE, Kay SA. Arabidopsis circadian clock protein, TOC1, is a DNA-binding transcription factor. Proceedings of the National Academy of Sciences of the United States of America. 2012;109(8):3167–72.

19. Huang W, Perez-Garcia P, Pokhilko A, Millar AJ, Antoshechkin I, Riechmann JL, et al. Mapping the core of the Arabidopsis circadian clock defines the network structure of the oscillator. Science. 2012;336(6077):75–9.

20. Nakamichi N, Kita M, Ito S, Yamashino T, Mizuno T. PSEUDO-RESPONSE REGULATORS, PRR9, PRR7 and PRR5, together play essential roles close to the circadian clock of Arabidopsis thaliana. Plant & cell physiology. 2005;46(5):686–98.

21. Nakamichi N, Kiba T, Henriques R, Mizuno T, Chua NH, Sakakibara H. PSEUDO-RESPONSE REGULATORS 9, 7, and 5 are transcriptional repressors in the Arabidopsis circadian clock. Plant Cell. 2010;22(3):594–605.

22. Nakamichi N, Kiba T, Kamioka M, Suzuki T, Yamashino T, Higashiyama T, et al. Transcriptional repressor PRR5 directly regulates clock-output pathways. Proceedings of the National Academy of Sciences of the United States of America. 2012;109(42):17123–8.

23. Elowitz MB, Leibler S. A synthetic oscillatory network of transcriptional regulators. Nature. 2000;403(6767):335–8.

24. Fogelmark K, Troein C. Rethinking Transcriptional Activation in the <italic>Arabidopsis</italic> Circadian Clock. PLoS Comput Biol. 2014;10(7):e1003705.

25. Joanito I, Chu J-W, Wu S-H, Hsu C-P. An incoherent feed-forward loop switches the Arabidopsis clock rapidly between two hysteretic states. Scientific reports. 2018;8(1):13944.

26. De Caluwé J, Xiao Q, Hermans C, Verbruggen N, Leloup J-C, Gonze D. A Compact Model for the Complex Plant Circadian Clock. Frontiers in Plant Science. 2016;7:74.

27. Wu J-F, Tsai H-L, Joanito I, Wu Y-C, Chang C-W, Li Y-H, et al. LWD–TCP complex activates the morning gene CCA1 in Arabidopsis. Nature Communications. 2016;7:13181.

28. Nusinow DA, Helfer A, Hamilton EE, King JJ, Imaizumi T, Schultz TF, et al. The ELF4-ELF3-LUX complex links the circadian clock to diurnal control of hypocotyl growth. Nature. 2011;475(7356):398–402.

29. Wang L, Fujiwara S, Somers DE. PRR5 regulates phosphorylation, nuclear import and subnuclear localization of TOC1 in the Arabidopsis circadian clock. The EMBO journal. 2010;29(11):1903–15.

30. Daniel X, Sugano S, Tobin EM. CK2 phosphorylation of CCA1 is necessary for its circadian oscillator function in <em>Arabidopsis</em>. Proceedings of the National Academy of Sciences of the United States of America. 2004;101(9):3292–7.

31. Mas P, Kim WY, Somers DE, Kay SA. Targeted degradation of TOC1 by ZTL modulates circadian function in Arabidopsis thaliana. Nature. 2003;426(6966):567–70.

32. Vierstra RD. The ubiquitin–26S proteasome system at the nexus of plant biology. Nature Reviews Molecular Cell Biology. 2009;10:385.

33. Shu K, Yang W. E3 Ubiquitin Ligases: Ubiquitous Actors in Plant Development and Abiotic Stress Responses. Plant and Cell Physiology. 2017;58(9):1461–76.

34. Kangisser S, Yakir E, Green RM. Proteasomal regulation of CIRCADIAN CLOCK ASSOCIATED 1 (CCA1) stability is part of the complex control of CCA1. Plant signaling & behavior. 2013;8(3):e23206.

35. Song HR, Carre IA. DET1 regulates the proteasomal degradation of LHY, a component of the Arabidopsis circadian clock. Plant molecular biology. 2005;57(5):761–71.

36. Park BS, Eo HJ, Jang IC, Kang HG, Song JT, Seo HS. Ubiquitination of LHY by SINAT5 regulates flowering time and is inhibited by DET1. Biochemical and biophysical research communications. 2010;398(2):242–6.

37. Ito S, Nakamichi N, Kiba T, Yamashino T, Mizuno T. Rhythmic and light-inducible appearance of clock-associated pseudo-response regulator protein PRR9 through programmed degradation in the dark in Arabidopsis thaliana. Plant Cell Physiol. 2007;48(11):1644–51.

38. Farre EM, Kay SA. PRR7 protein levels are regulated by light and the circadian clock in Arabidopsis. The Plant journal : for cell and molecular biology. 2007;52(3):548–60.

39. Kiba T, Henriques R, Sakakibara H, Chua NH. Targeted degradation of PSEUDO-RESPONSE REGULATOR5 by an SCFZTL complex regulates clock function and photomorphogenesis in Arabidopsis thaliana. Plant Cell. 2007;19(8):2516–30.

40. David KM, Armbruster U, Tama N, Putterill J. Arabidopsis GIGANTEA protein is post-transcriptionally regulated by light and dark. FEBS letters. 2006;580(5):1193–7.

41. Yu JW, Rubio V, Lee NY, Bai S, Lee SY, Kim SS, et al. COP1 and ELF3 control circadian function and photoperiodic flowering by regulating GI stability. Mol Cell. 2008;32(5):617–30.

42. Takahashi JS. Transcriptional architecture of the mammalian circadian clock. Nature Reviews Genetics. 2016;18:164.

43. Siepka SM, Yoo S-H, Park J, Song W, Kumar V, Hu Y, et al. Circadian Mutant Overtime Reveals F-box Protein FBXL3 Regulation of Cryptochrome and Period Gene Expression. Cell. 2007;129(5):1011–23.

44. Godinho SIH, Maywood ES, Shaw L, Tucci V, Barnard AR, Busino L, et al. The After-Hours Mutant Reveals a Role for Fbxl3 in Determining Mammalian Circadian Period. Science. 2007;316(5826):897–900.

45. Busino L, Bassermann F, Maiolica A, Lee C, Nolan PM, Godinho SIH, et al. SCF<sup>Fbxl3</sup> Controls the Oscillation of the Circadian Clock by Directing the Degradation of Cryptochrome Proteins. Science. 2007;316(5826):900–4.

46. Hirano A, Yumimoto K, Tsunematsu R, Matsumoto M, Oyama M, Kozuka-Hata H, et al. FBXL21 Regulates Oscillation of the Circadian Clock through Ubiquitination and Stabilization of Cryptochromes. Cell. 2013;152(5):1106–18.

47. Yoo S-H, Mohawk Jennifer A, Siepka Sandra M, Shan Y, Huh Seong K, Hong H-K, et al. Competing E3 Ubiquitin Ligases Govern Circadian Periodicity by Degradation of CRY in Nucleus and Cytoplasm. Cell. 2013;152(5):1091–105.

48. Pokhilko A, Fernandez AP, Edwards KD, Southern MM, Halliday KJ, Millar AJ. The clock gene circuit in Arabidopsis includes a repressilator with additional feedback loops. Mol Syst Biol. 2012;8:574.

49. Pett JP, Korenčič A, Wesener F, Kramer A, Herzel H. Feedback Loops of the Mammalian Circadian Clock Constitute Repressilator. PLOS Computational Biology. 2016;12(12):e1005266.

50. Tsai TY, Choi YS, Ma W, Pomerening JR, Tang C, Ferrell JE, Jr. Robust, tunable biological oscillations from interlinked positive and negative feedback loops. Science. 2008;321(5885):126–9.

51. Strelkowa N, Barahona M. Switchable genetic oscillator operating in quasi-stable mode. Journal of The Royal Society Interface. 2010;7(48):1071–82.

52. Perez-Carrasco R, Barnes CP, Schaerli Y, Isalan M, Briscoe J, Page KM. Combining a Toggle Switch and a Repressilator within the AC-DC Circuit Generates Distinct Dynamical Behaviors. Cell Syst. 2018.

53. Gillespie DT. A general method for numerically simulating the stochastic time evolution of coupled chemical reactions. Journal of Computational Physics. 1976;22(4):403–34.

54. Gillespie DT. Exact stochastic simulation of coupled chemical reactions. The Journal of Physical Chemistry. 1977;81(25):2340–61.

55. Wang B, Kitney RI, Joly N, Buck M. Engineering modular and orthogonal genetic logic gates for robust digital-like synthetic biology. Nature Communications. 2011;2:508.

56. Mertens N, Remaut E, Fiers W. Tight Transcriptional Control Mechanism Ensures Stable High-Level Expression from T7 Promoter-Based Expression Plasmids. Bio/Technology. 1995;13:175.

57. Lee TS, Krupa RA, Zhang F, Hajimorad M, Holtz WJ, Prasad N, et al. BglBrick vectors and datasheets: A synthetic biology platform for gene expression. Journal of Biological Engineering. 2011;5(1):12.

58. Huang HH, Camsund D, Lindblad P, Heidorn T. Design and characterization of molecular tools for a Synthetic Biology approach towards developing cyanobacterial biotechnology. Nucleic acids research. 2010;38(8):2577–93.

59. Yanai I, Korbel JO, Boue S, McWeeney SK, Bork P, Lercher MJ. Similar gene expression profiles do not imply similar tissue functions. Trends in Genetics. 2006;22(3):132–8.

60. Ochab-Marcinek A, Tabaka M. Transcriptional leakage versus noise: a simple mechanism of conversion between binary and graded response in autoregulated genes. Physical review E, Statistical, nonlinear, and soft matter physics. 2015;91(1):012704.

61. Barkai N, Leibler S. Biological rhythms: Circadian clocks limited by noise. Nature. 2000;403(6767):267–8.

62. Vilar JMG, Kueh HY, Barkai N, Leibler S. Mechanisms of noise-resistance in genetic oscillators. Proceedings of the National Academy of Sciences. 2002;99(9):5988–92.

63. Avendaño MS, Leidy C, Pedraza JM. Tuning the range and stability of multiple phenotypic states with coupled positive–negative feedback loops. Nature Communications. 2013;4:2605.

64. Hornung G, Barkai N. Noise Propagation and Signaling Sensitivity in Biological Networks: A Role for Positive Feedback. PLOS Computational Biology. 2008;4(1):e8.

65. Kim WY, Geng R, Somers DE. Circadian phase-specific degradation of the F-box protein ZTL is mediated by the proteasome. Proceedings of the National Academy of Sciences of the United States of America. 2003;100(8):4933–8.

66. Kim WY, Fujiwara S, Suh SS, Kim J, Kim Y, Han L, et al. ZEITLUPE is a circadian photoreceptor stabilized by GIGANTEA in blue light. Nature. 2007;449(7160):356–60.

67. Jin J, Tian F, Yang D-C, Meng Y-Q, Kong L, Luo J, et al. PlantTFDB 4.0: toward a central hub for transcription factors and regulatory interactions in plants. Nucleic acids research. 2016;45(D1):D1040–D5.

68. Chow C-N, Lee T-Y, Hung Y-C, Li G-Z, Tseng K-C, Liu Y-H, et al. PlantPAN3.0: a new and updated resource for reconstructing transcriptional regulatory networks from ChIP-seq experiments in plants. Nucleic acids research. 2018;47(D1):D1155–D63.

69. Huang H, Alvarez S, Bindbeutel R, Shen Z, Naldrett MJ, Evans BS, et al. Identification of Evening Complex Associated Proteins in <em>Arabidopsis</em> by Affinity Purification and Mass Spectrometry. Molecular & Cellular Proteomics. 2016;15(1):201–17.

70. Del Punta K, Leinders-Zufall T, Rodriguez I, Jukam D, Wysocki CJ, Ogawa S, et al. Deficient pheromone responses in mice lacking a cluster of vomeronasal receptor genes. Nature. 2002;419(6902):70–4.

71. Hazen SP, Schultz TF, Pruneda-Paz JL, Borevitz JO, Ecker JR, Kay SA. LUX ARRHYTHMO encodes a Myb domain protein essential for circadian rhythms. Proceedings of the National Academy of Sciences of the United States of America. 2005;102(29):10387–92.

72. Dixon LE, Knox K, Kozma-Bognar L, Southern MM, Pokhilko A, Millar AJ. Temporal repression of core circadian genes is mediated through EARLY FLOWERING 3 in Arabidopsis. Current biology : CB. 2011;21(2):120–5.

73. Nagel DH, Doherty CJ, Pruneda-Paz JL, Schmitz RJ, Ecker JR, Kay SA. Genome-wide identification of CCA1 targets uncovers an expanded clock network in Arabidopsis. Proceedings of the National Academy of Sciences. 2015;112(34):E4802–E10.

74. Adams S, Grundy J, Veflingstad SR, Dyer NP, Hannah MA, Ott S, et al. Circadian control of abscisic acid biosynthesis and signalling pathways revealed by genome-wide analysis of LHY binding targets. New Phytologist. 2018;220(3):893–907.

75. Liu TL, Newton L, Liu M-J, Shiu S-H, Farré EM. A G-Box-Like Motif Is Necessary for Transcriptional Regulation by Circadian Pseudo-Response Regulators in Arabidopsis. Plant physiology. 2016;170(1):528–39.

76. Ito S, Nakamichi N, Matsushika A, Fujimori T, Yamashino T, Mizuno T. Molecular dissection of the promoter of the light-induced and circadian-controlled APRR9 gene encoding a clock-associated component of Arabidopsis thaliana. Bioscience, biotechnology, and biochemistry. 2005;69(2):382–90.

77. Stachel SE, Zambryski PC. virA and virG control the plant-induced activation of the T-DNA transfer process of A. tumefaciens. Cell. 1986;46(3):325–33.

78. Yamamoto A, Iwahashi M, Yanofsky MF, Nester EW, Takebe I, Machida Y. The promoter proximal region in the virD locus of Agrobacterium tumefaciens is necessary for the plant-inducible circularization of T-DNA. Molecular and General Genetics MGG. 1987;206(1):174–7.

79. DiGiuseppe PA, Silhavy TJ. Signal Detection and Target Gene Induction by the CpxRA Two-Component System. Journal of Bacteriology. 2003;185(8):2432–40.

80. Kyrkanides S, Miller J-nH, Bowers WJ, Federoff HJ. Transcriptional and posttranslational regulation of cre recombinase by ru486 as the basis for an enhanced inducible expression system. Molecular Therapy. 2003;8(5):790–5.

81. Tigges M, Marquez-Lago TT, Stelling J, Fussenegger M. A tunable synthetic mammalian oscillator. Nature. 2009;457:309.

82. Danino T, Mondragón-Palomino O, Tsimring L, Hasty J. A synchronized quorum of genetic clocks. Nature. 2010;463:326.

83. Deans TL, Cantor CR, Collins JJ. A Tunable Genetic Switch Based on RNAi and Repressor Proteins for Regulating Gene Expression in Mammalian Cells. Cell. 2007;130(2):363–72.

84. Avery SV. Cell individuality: the bistability of competence development. Trends in Microbiology. 2005;13(10):459–62.

85. Ingolia NT, Murray AW. Positive-Feedback Loops as a Flexible Biological Module. Current Biology. 2007;17(8):668–77.

86. Alon U. An Introduction to Systems Biology: Design Principles of Biological Circuits: Taylor & Francis; 2006.

87. Ferrell JE, Jr., Ha SH. Ultrasensitivity part II: multisite phosphorylation, stoichiometric inhibitors, and positive feedback. Trends in biochemical sciences. 2014;39(11):556–69.

88. Fujiwara S, Wang L, Han L, Suh SS, Salome PA, McClung CR, et al. Post-translational regulation of the Arabidopsis circadian clock through selective proteolysis and phosphorylation of pseudo-response regulator proteins. The Journal of biological chemistry. 2008;283(34):23073–83.

89. Yakir E, Hilman D, Kron I, Hassidim M, Melamed-Book N, Green RM. Posttranslational regulation of CIRCADIAN CLOCK ASSOCIATED1 in the circadian oscillator of Arabidopsis. Plant physiology. 2009;150(2):844–57.

90. Pokhilko A, Hodge SK, Stratford K, Knox K, Edwards KD, Thomson AW, et al. Data assimilation constrains new connections and components in a complex, eukaryotic circadian clock model. Mol Syst Biol. 2010;6:416.

91. Cao Y, Gillespie DT, Petzold LR. Efficient step size selection for the tau-leaping simulation method. The Journal of chemical physics. 2006;124(4):044109.

92. Golding I, Paulsson J, Zawilski SM, Cox EC. Real-time kinetics of gene activity in individual bacteria. Cell. 2005;123(6):1025–36.

93. Yu J, Xiao J, Ren X, Lao K, Xie XS. Probing gene expression in live cells, one protein molecule at a time. Science. 2006;311(5767):1600–3.

94. Yan C-CS, Chepyala SR, Yen C-M, Hsu C-P. Efficient and flexible implementation of Langevin simulation for gene burst production. Scientific reports. 2017;7(1):16851.

95. van Dorp M, Lannoo B, Carlon E. Generation of oscillating gene regulatory network motifs. Physical review E, Statistical, nonlinear, and soft matter physics. 2013;88(1):012722.

96. Otero-Muras I, Banga JR. Design Principles of Biological Oscillators through Optimization: Forward and Reverse Analysis. PLOS ONE. 2016;11(12):e0166867.

